# Mapping the canine microbiome: Insights from the Dog Aging Project

**DOI:** 10.1101/2024.12.02.625632

**Authors:** Tal Bamberger, Efrat Muller, Yadid M. Algavi, Ashlee Greenier, Christine Adjangba, Elizabeth Slikas, Layla Brassington, Blaise Mariner, Brianah McCoy, Benjamin R. Harrison, Maria Partida-Aguilar, Abbey Marye, Adam Harris, Emily Rout, DAP Consortium, Anne Avery, Daniel E.L. Promislow, Noah Snyder-Mackler, Elhanan Borenstein

**Author notes:** Equal contribution (Tal Bamberger, Efrat Muller, Yadid M. Algavi). Corresponding author: **Elhanan Borenstein,**.

## Abstract

Companion dogs (*Canis lupus familiaris*) offer a unique model for studying the gut microbiome and its relation to aging due to their cohabitation with humans, sharing similar environments, diets, and healthcare practices. Here, we present the Dog Aging Project (DAP) Precision cohort, the largest population-wide study of the canine gut microbiome to date. This cohort encompasses over 900 dogs of diverse breeds, environments, and demographics living across the United States. Coupling fecal shotgun metagenomic sequencing with comprehensive phenotypic and environmental surveys and clinical lab tests, we explore the intricate relationships between microbiome composition, aging, and key factors such as diet, health, and living conditions. Our analyses identify various factors associated with microbiome composition. In addition, we find a gradual shift in microbiome composition with age, which allows us to develop a novel metagenomics-based “clock” to predict biological aging based on microbial signatures. Overall, these findings provide an unprecedented and detailed understanding of the role the gut microbiome plays in our four-legged companions, offering both potential applications in veterinary medicine and an exciting model for aging research.

## Introduction

The past two decades have witnessed a surge of interest in host-associated microbiomes, the complex communities of microorganisms that inhabit various ecological niches on and within host organisms. Extensive research, primarily focused on the human microbiome, has revealed the profound impact that the microbiome exerts on the host’s health, influencing key aspects of digestion, immunity, aging, and even behavior^1^. In turn, lifestyle choices, diet, host genetics, and pharmaceutical interventions have emerged as crucial factors shaping the function and composition of the microbiome^2,3^. Of the various host processes associated with and impacting the microbiome, perhaps the most intriguing and tightly linked is aging. Indeed, a growing body of evidence established a strong association between the microbiome and aging^4–6^, demonstrating clear age-associated shifts in the microbiome and mechanisms by which the microbiome may impact host aging^7,8^. The microbiome is also associated with multiple age-related clinical conditions in humans, including frailty, colorectal carcinoma, and atherosclerotic disease^9,10^.

As with many physiological processes, the study of aging and its relationship to the gut microbiome cannot rely purely on human-based studies, and calls for a robust, effective, and clinically-relevant model system. The choice of such a model system is especially challenging in microbiome research given the dynamic nature of the microbiome and its interaction with the host environment and lifestyle. Murine models, for example, widely used in biomedical research in general and in microbiome studies specifically, allow for controlled experimental conditions that make it possible to assess causality in complex host-microbiota interactions. Nonetheless, murine models do not share many of the environmental and lifestyle factors that shape the human gut microbiome, and lack the genetic diversity of humans^11^. Similarly, existing microbiome studies in free-living mammals often reflect a trade-off between observing animals in their natural habitats and studying them in controlled environments. Research on wild primates such as baboons^12^, macaques^13^, and gelada^14^, for example, offers valuable insights into microbial ecology, diversity, and activity in naturally occurring host-associated microbiomes, but is complicated by many confounding factors such as variations in diet and health conditions. On the other hand, controlled settings, such as those of cows in dairy farms^15^ or chickens in hen houses^16^, provide improved control and repeatability, yet fail to replicate the complexities and multifaceted influences of human lifestyles and environments.

Companion dogs (*Canis lupus familiaris*), however, present a unique opportunity to bridge this gap, offering an invaluable model system for studying host-microbiome-environment interactions. Indeed, as integral members of human households, dogs share not only our living spaces but also key lifestyle factors, demographics, and environmental exposures^17^. Indeed, previous studies of the canine microbiome have revealed important links between the microbiome and various host phenotypes or interventions, mimicking associations that were also previously indicated in humans. These include, for example, response to dietary change^18,19^, carriage of antibiotic resistance genes^20,21^ and disease-specific alterations in inflammatory bowel disease^22^ and atopic dermatitis^23^. Moreover, a recent study, analyzing over 2,000 canine stool microbiomes, revealed that certain microbial species are shared between dogs and humans^20^. Additionally, companion dogs benefit from advanced healthcare access, rendering any observation that could inform canine healthcare a valuable finding in and of itself.

In the context of aging, several studies in dogs have demonstrated intriguing age-associated shifts in the gut microbiome composition. Fernández-Pinteño *et al.*^24^, for example, recently analyzed 16S sequenced fecal samples from 106 dogs of diverse breeds, and reported that while no differences in alpha diversity were identified between age groups, several specific bacterial families were significantly more abundant in senior dogs compared to younger age groups. In another study, Mizukami *et al.*^25^ identified a decrease in microbial diversity with age, in a cohort of 43 purebred Shiba Inu dogs. Moreover, Kubinyi *et al.*^26^ assessed 29 pet dogs, finding an association between the composition of the gut microbiome and both age and memory. Importantly, however, these studies (and others) are limited in scale, focus on specific breeds, or employ a cross-sectional study design, all of which could restrict our ability to capture the full diversity of companion dogs and their gut microbiomes, ultimately falling short of providing a large-scale, comprehensive mapping of the various determinants of the canine gut microbiome.

To overcome these limitations and gain deeper insights into how diverse, real-world environments and lifestyle choices influence host-microbiome interactions, here we use data from the Dog Aging Project (DAP), the largest population-wide longitudinal study of companion dogs to date^27^. The DAP follows over 50,000 companion dogs across the United States, collecting detailed genetic, clinical, environmental, and lifestyle data to investigate the biological and environmental determinants of healthy aging. Specifically, as part of this project, we have enrolled nearly a thousand dogs, representing a variety of breeds, lifestyles, and demographics across the United States, into the Dog Aging Project Precision Cohort. This cohort includes a subset of the broader DAP population with comprehensive omics data, including extensive phenotypic and environmental survey data, clinical lab tests, and fecal microbiome data using shotgun metagenomic sequencing. In this study, focusing specifically on the dog gut microbiome, we characterize the bacterial community’s structure and function, and explore associations between the microbiome profiles and various dietary, environmental, and clinical factors. We identify significant associations between age and the composition of the gut microbial community, suggesting how the aging process may shape the canine microbiome. Our analyses also reveal strong associations between microbes and clinical lab tests. Overall, this analysis provides valuable insights into the complex interplay between dogs and their microbial inhabitants, with potential implications for veterinary medicine and human health.

## Results

### The Dog Aging Project (DAP) Precision Cohort

In this study, we analyzed gut microbiome profiles of dogs that took part in the first-year Precision Cohort of the Dog Aging Project (DAP)^27^. These dogs are surveyed for environmental, physiological, and molecular factors, including gut microbiome composition, as well as other molecular omics analyzed and reported elsewhere^28–31^. Overall, 976 dogs were enrolled in this cohort, of which 922 were successfully sampled and processed for gut microbiome profiles and included in our analysis. This large-scale dataset provides a comprehensive representation of canine demographics across the United States (Figure 1A). Notably, the dogs analyzed here included a balanced sex distribution, with 444 females and 478 males, 84% of whom (from both sexes) were sterilized. The cohort captured most of the canine lifespan (with age ranging from 8 months to 18 years, with a median of 4.6 years; Figure 1B-C), and included diverse breeds, with the most common being Golden Retriever, German Shepherd, and Labrador Retriever (Figure 1D), and 48% of the dogs being mixed-breed. The cohort also represented the full range of American Veterinary Medical Association (AVMA)-designated breed size classes and life stages (Figure 1E). Owner-reported health status indicated that most of the dogs were in good health at the time of enrollment (Figure 1F). Overall, this cohort thus reflected a wide and diverse range of companion dogs, including a comprehensive profiling of each cohort member, and offered an invaluable opportunity to study the factors linking the canine gut microbiome to canine aging.

**Figure 1:**
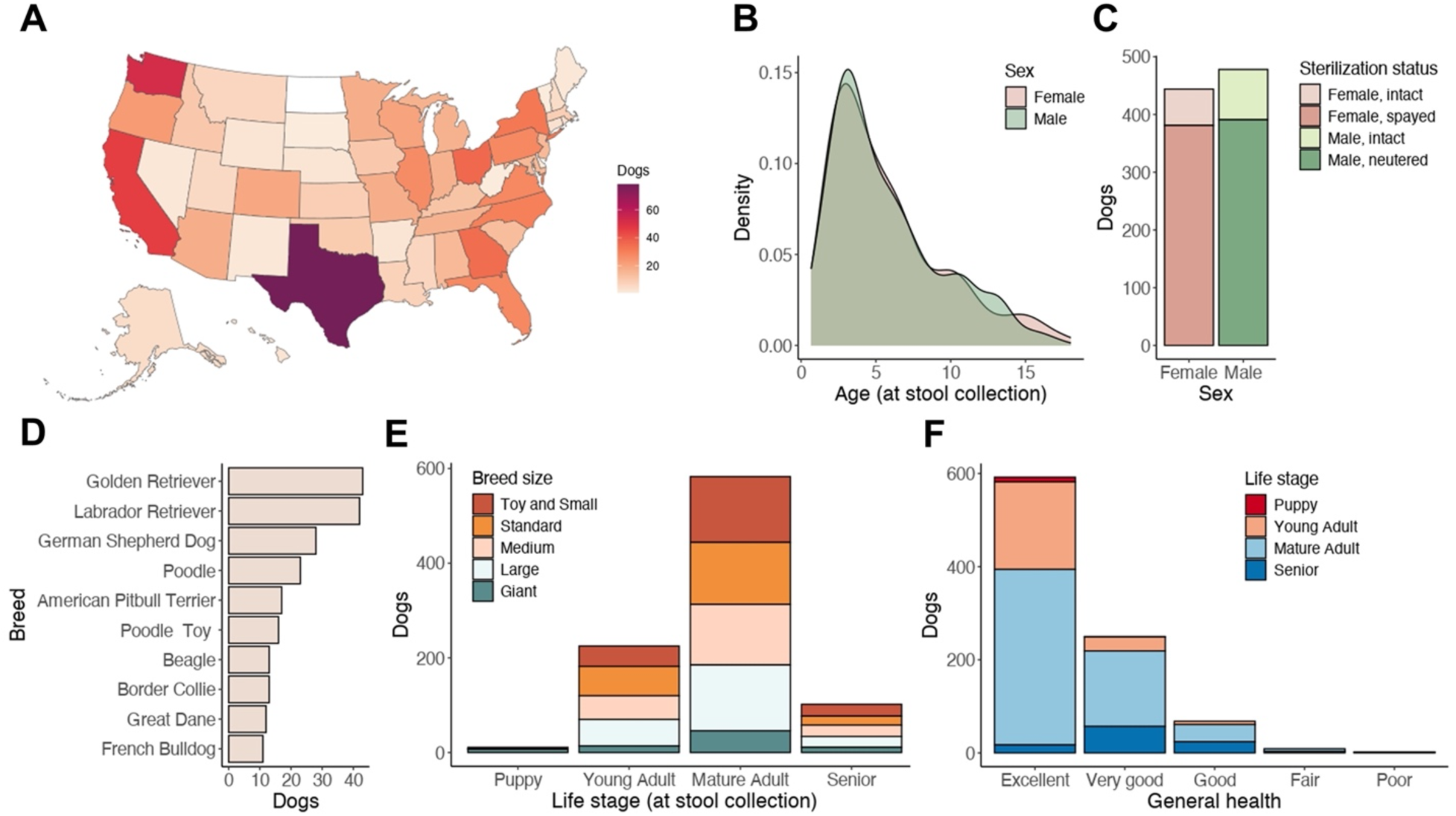
The Dog Aging Project Precision Cohort overview. **(A)** A map showing the geographic distribution of the 922 dogs across the United States. **(B)** Density plot illustrating the age distribution of the dogs included in our study, colored by sex. **(C)** A bar plot showing the number of dogs of each sex, colored by sterilization status. **(D)** A bar plot presenting the number of dogs in the ten most abundant breeds (among purebred dogs). **(E)** A bar plot representing the number of dogs of each breed size, across different life stage categories. Colors indicate breed size classes. **(F)** A bar plot showing the distribution of owner-reported health status of the dogs, across different life stage categories.

### Composition of the gut microbiome in DAP dogs and its correlates

To characterize the composition of the gut microbiome of DAP dogs, we obtained fecal samples from 922 dogs, performed shotgun metagenomic sequencing of these samples, and processed the obtained sequencing reads to construct the taxonomic and functional profiles of each dog’s microbiome (see Methods). The most abundant bacterial genera observed in the DAP dogs’ fecal microbiomes, based on GTDB taxonomy^32^ were *Prevotella* (average relative abundance 28.1%), *Phocaeicola* (26.1%; although, in NCBI, many *Phocaeicola* strains are classified as *Bacteroides*), *Bacteroides* (5.6%), and *Alloprevotella* (4.3%) from the Bacteroidota phylum, *Fusobacterium_A* (6%) from the Fusobacteriota phylum, and *Escherichia* (3.3%) and *Sutterella* (3.2%) from the Pseudomonadota (Proteobacteria) phylum. All of the above genera were also highly prevalent, present in >90% of the samples (Figure 2A; Supplementary Table S1). Notably, these findings are in line with previous reports of the dog gut microbiome that were based on 16S rRNA gene amplicon sequencing^26^. At the species level, *Phocaeicola sp900546645* (8.3%), *Prevotella sp900551275* (7.8%; previously referred to as *Segatella copri*), *Phocaeicola vulgatus* (5.6%), *Prevotella copri* (5.3%), *Prevotella sp015074785* (4.3%), and *Phocaeicola coprocola* (4.1%) were the most abundant (Supplementary Table S1). To validate that DAP microbiome profiles are consistent with those obtained in previous dog studies, we reprocessed data from the largest dog cohort that had fecal shotgun metagenomic data to date, including 32 Beagles and 32 Labradors^18^. We confirmed that DAP taxonomic profiles were highly similar to those observed in this previous cohort, with significant correlations between the average relative abundances of species (Spearman ρ=0.69, p<10^-10^) and genera (Spearman ρ=0.55, p<10^-8^) in the two cohorts (Figure 2B; Supplementary Figure S1). We further compared average species’ abundances in DAP, computed using a reference-based approach and GTDB taxonomy, to those computed in a recent study by Branck et al.^20^, which used a different reference database and further incorporated newly assembled genomes. We found that despite these differences in processing, the average relative abundances of gut species in DAP were significantly and highly correlated with those obtained in that study (Supplementary Figure S2, Spearman ρ=0.62, p<10^-5^; Methods).

**Figure 2:**
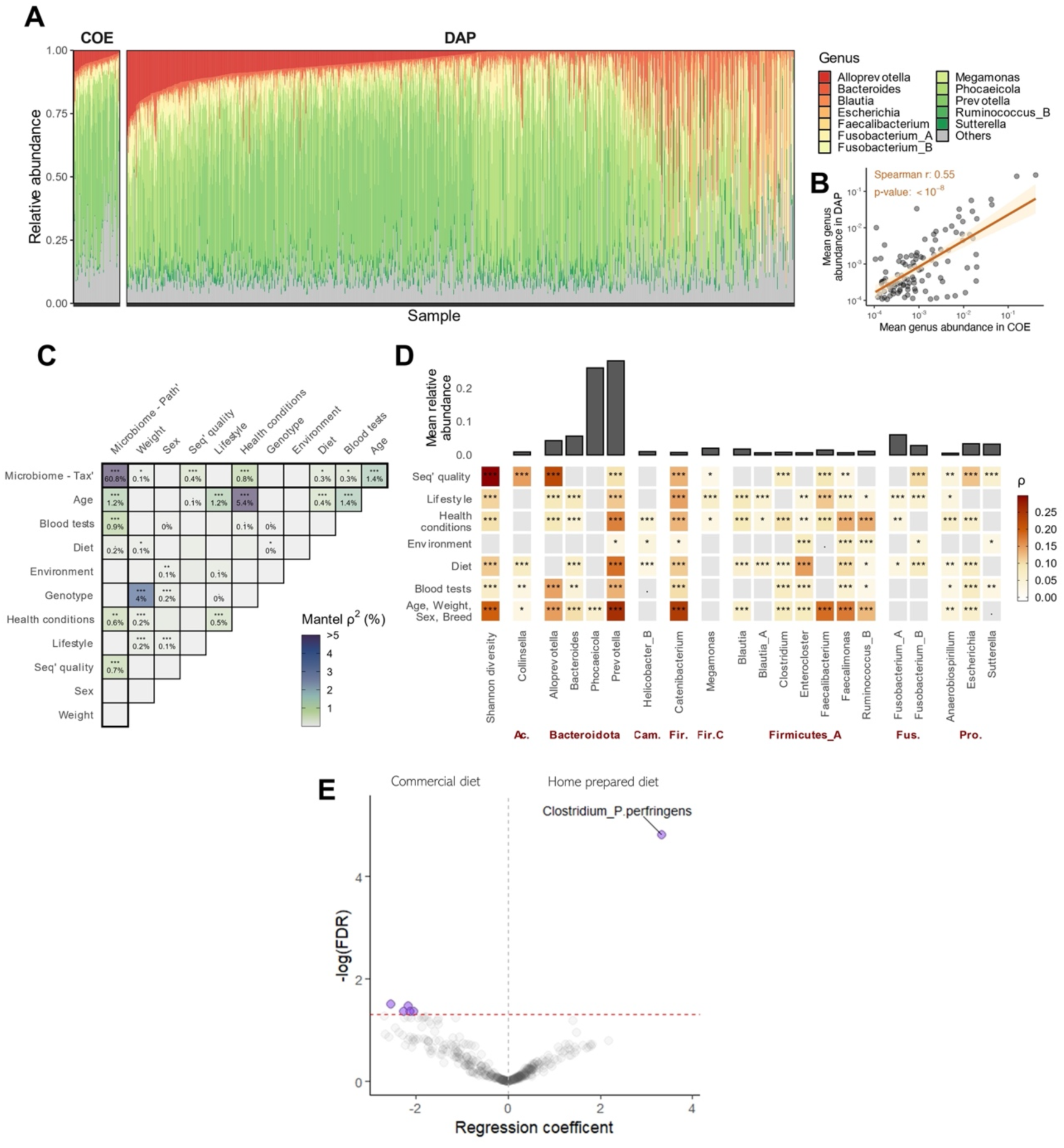
The gut microbiome of study dogs and key factors associated with it. **(A)** Genus-level bacterial compositions of fecal samples from 922 Precision 1 DAP samples (“DAP”), along with 64 fecal samples from Coehlo et al. (“COE”). Genera for which the average relative abundance was below 1% were grouped into an “Others” category and colored in grey. **(B)** A comparison of average genera abundances between the two cohorts (“DAP” and “COE” as above). Each point in the scatter point represents a single genus, for which its average relative abundance in each dataset was calculated. Regression line and correlation statistics are presented in orange. **(C)** Pairwise Mantel tests to estimate the associations between each pair of measurement types. Tiles are colored by the squared Mantel’s statistic (Spearman’s rho), roughly equivalent to shared variance. Stars indicate FDR-corrected p-value significance (. ≤ 0.1; * ≤ 0.05; ** ≤ 0.01; *** ≤ 0.001). See Supplementary Table S3. Tax’: Taxonomy, Path’: Pathways, Seq’: Sequencing. **(D)** Random forest regression models were used to predict either Shannon diversity or relative abundances of the top 20 most abundant genera (x-axis), based on different data types (y-axis). Models were evaluated using 10 repeats of 5-fold cross validation, and tiles are colored based on the Spearman correlation between predicted and true values, averaged over folds. Stars indicate FDR-corrected p-value significance of the Spearman correlation, as noted above. Top panel presents average relative abundances of each genus. **(E)** A volcano plot illustrating the differential abundance of microbial species between dogs fed home-prepared diets and those fed commercial diets. Each point represents a microbial species, with the x-axis showing the effect size (mixed effects model coefficient) and the y-axis indicating the -log10 adjusted p-value (FDR).

Next, we sought to identify correlates of the obtained taxonomic and functional fecal microbiome profiles, exploring broad associations between these profiles and multiple demographic, lifestyle, and health-related factors of the DAP dogs, available from owner-reported surveys and clinical lab tests (Supplementary Table S2). Using sample distance matrices computed for each data type, we found that dog age, blood test results, diet, and health conditions, were all significantly associated with both the taxonomic and functional profiles of the microbiome (Figure 2C; Supplementary Figure 3A; Supplementary Table S3; FDR<0.05, Mantel and Procrustes tests; Methods). Furthermore, dogs’ age was significantly associated with taxonomic composition, even when controlling for all other factors (Supplementary Figure 3B; Supplementary Table S4; Methods). Measures related to sequencing quality were also significantly associated with microbiome composition and were therefore included as covariates in all downstream analyses.

To further explore whether such factors are associated with general community-level shifts or can explain variation in specific bacterial genera, we tested if they could be used to predict the relative abundances of specific genera. Training random forest regression models with cross-validation to evaluate this, we were able to successfully predict the abundances of multiple different genera using various data types (Figure 2D; Supplementary Table S5). Overall, these results suggest prevalent and widespread associations between the dog gut microbiome and a variety of demographic, lifestyle, and health-related factors.

Next, we used information from the DAP owner surveys to explore how different dietary choices are associated with the canine gut microbiome at a population-wide scale. We observed significant differences in the microbiomes of dogs based on their primary diet (FDR=0.001, Bray-Curtis PERMANOVA), with 34 species, 9 genera, and 13 KEGG modules that were differentially abundant among dietary groups (such as commercially prepared dry food, refrigerated or frozen raw food, home prepared raw diet, or home prepared cooked diet; Supplementary Table S6; FDR<0.05, mixed effects model). Grouping dogs into two main dietary categories, namely home-prepared diets vs. commercial-diets, we further identified 5 microbial species whose relative abundances were significantly lower in the microbiomes of dogs that were fed a home-prepared diet, and 1 species, *Clostridium perfringens*, that was significantly higher in these dogs. *Clostridium perfringens* is a known pathogen that can cause diarrhea in dogs, and its overabundance in dogs fed home-prepared diets may thus highlight a potential risk associated with home-prepared meals^33^. Additionally, grain-free diets, which often include alternative carbohydrate and fiber sources such as peas, lentils, or potatoes, were also significantly associated with the microbiome profile (FDR=0.003, Bray-Curtis PERMANOVA), with 21 differentially abundant species, genera, or KEGG modules (Supplementary Table S6; FDR<0.05, mixed effects model).

While food supplements such as antioxidants, vitamins, probiotics, omega-3 fatty acids, chondroitin sulfate, dietary fibers, or glucosamine supplements were not significantly associated with the microbial community structure (FDR>0.05, PERMANOVA), coprophagy (feces eating) emerged as a significant correlate (FDR=0.002, Bray-Curtis PERMANOVA). Interestingly, coprophagy was associated with higher Shannon diversity (FDR=0.0002, Wilcoxon test), potentially due to a higher influx of environmental microbial species. This habit was also associated with higher abundance of 10 bacterial species and 5 genera (Supplementary Table S6; FDR<0.05, mixed effects model). Indeed, similar findings have been reported in rodents, rabbits, and birds, where coprophagy significantly influences gut microbiome composition^34–36^. Overall, these findings underscore the complexity of dietary influences on the canine microbiome and highlight the importance of considering both the type of diet and specific behaviors, such as coprophagy, in understanding microbiome composition.

### Association of the gut microbiome with age and the construction of a metagenomics-based clock

We next focused on investigating the association between dog age and the gut microbiome. We found that the microbiome alpha diversity (*i.e.*, the diversity of species within each dog’s gut microbiome), as measured by the Shannon index, significantly decreased with dog age, even when controlling for dogs’ health status, sex, weight, breed size, and technical covariates (Figure 3A; p=0.0002, mixed effects models). Moreover, significant differences were observed when comparing the mean distance of each dog to a randomly sampled reference group of young adult dogs (Figure 3B; FDR<0.0001). This was also consistent with observed microbiome composition differences between age groups, as measured by permutational multivariate ANOVA (PERMANOVA; Figure 3C; p=0.0001). Overall, these findings suggest that older age in dogs is associated with a less diverse gut microbiome and with shifts in microbiome composition.

**Figure 3:**
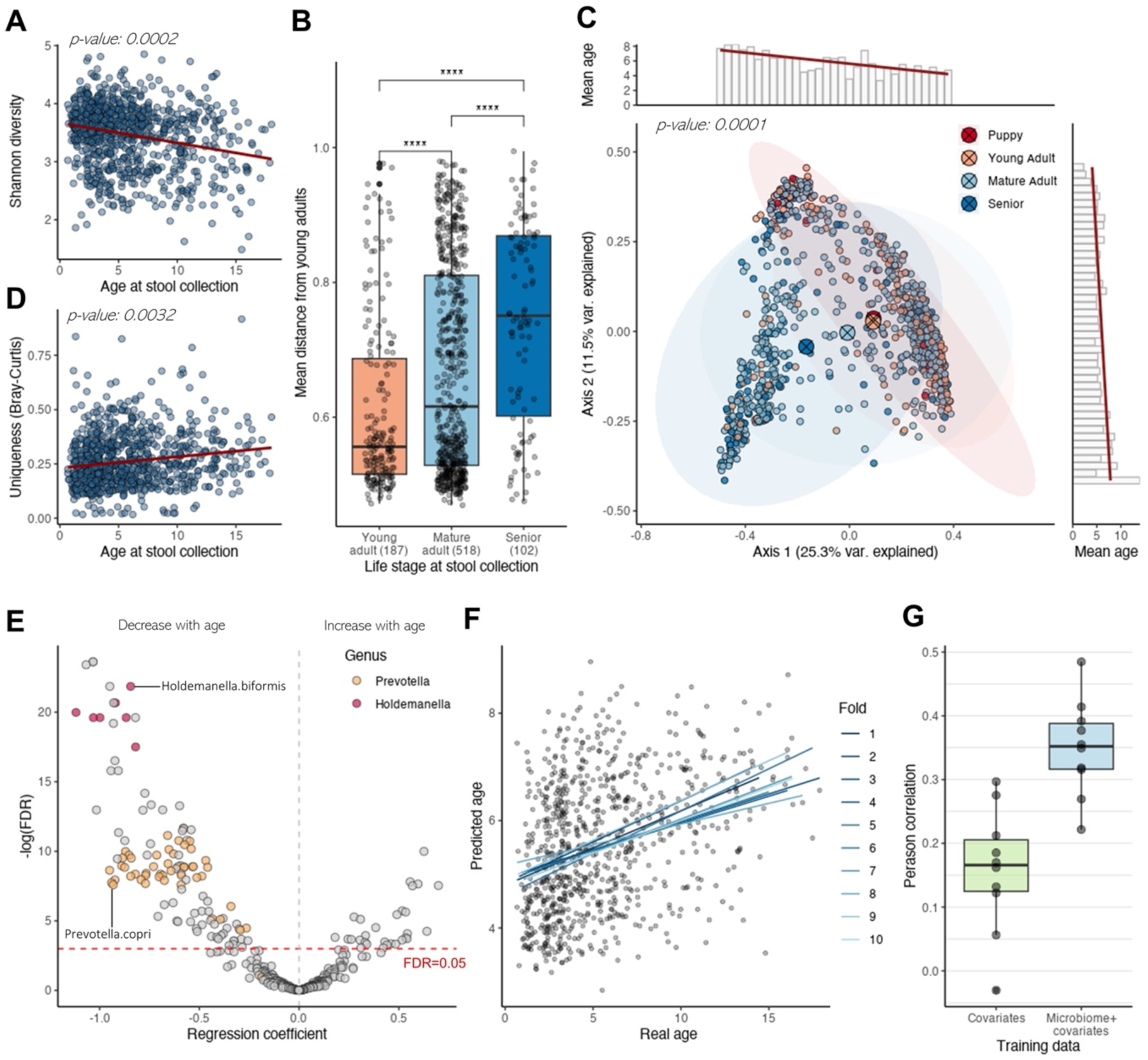
Microbiome associations with age. **(A)** Alpha Diversity (Shannon Diversity) of the microbiome association with age, as estimated by mixed effects modeling. **(B)** Boxplot of the mean distances of dogs in three age categories to a randomly sampled reference group of young adult dogs. Each dot represents the mean distance for an individual dog to all the dogs in the reference group, highlighting significant differences in the distribution of mean distances across the age groups. Statistical significance was determined using the Wilcoxon test, with significance levels denoted by asterisks, after FDR correction (* FDR < 0.05, ** FDR < 0.01, *** FDR < 0.001, **** FDR < 0.0001). **(C)** Principal Coordinate Analysis (PCoA) with Bray-Curtis distances, color-coded by life stage categories. Statistical significance was assessed using PERMANOVA (Permutational Multivariate Analysis of Variance). The bar plots at the margins describe the mean age along the first two PCoA axes, calculated as the mean age of the dogs in each bin along the respective axis after dividing the axis into bins. **(D)** Gut microbiome uniqueness (calculated using the species-level Bray-Curtis dissimilarity metric) association with age. Significance was evaluated via mixed effects modeling. **(E)** Volcano plot illustrating the differential abundance of species related to age, as calculated by MaAsLin2. Colors highlight two genus-level taxa: Prevotella and Holdemanella. p-values have been adjusted using FDR correction. **(F)** Correlation between real age and predicted age based on metagenomic data, analyzed using 10-fold cross-validation. Each line represents the correlation from a different fold. **(G)** Boxplot displaying the correlations for each fold when training models with both microbiome and covariates, compared to a baseline model trained only with covariates.

We additionally evaluated microbiome “uniqueness”, a trait describing how distinct an individual’s microbiome composition is compared to others within a given cohort, a metric recently reported to increase with healthy aging in humans^4^. We found that a dog’s gut microbiome uniqueness was indeed positively associated with age, controlling for all previously mentioned covariates (Figure 3D; p=0.0032, mixed effects models; see Methods). Furthermore, this trend was also significant when controlling for Shannon diversity (p=0.0113), following Wilmanski *et al.*^4^. Interestingly, when we stratified dogs into two health categories: the healthiest dogs and all others, this association remained significant for the healthiest group (p=0.0036), but was no longer significant for the other group (p=0.908). These findings mirror observations in humans, where uniqueness was characteristic of “healthy” aging only^4^ (although, this observation may also be due to the relatively small numbers of dogs in our cohort with poorer owner-reported health status; Figure 1F).

Next, we set out to identify specific microbiome taxa associated with age (see Methods). Our analysis, using a mixed effects model that controlled for dogs’ health status, sex, weight, breed size, and technical covariates, revealed significant associations between age and the abundance of multiple microbial taxa, as well as the abundance of functional modules and pathways. Specifically, we identified 147 species, 46 genera, and 7 phyla, alongside 59 KEGG pathways and 92 KEGG modules, with significant associations with dog age (Figure 3E; Supplementary Tables S7, S8, S9, S10, S11; FDR<0.05). Notable among these taxa were *Prevotella* and *Holdemanella*, which were significantly less abundant in older dogs at both the genus and the species levels, with 48 and 7 corresponding species, respectively. These findings are in line with previous observations concerning the link between these taxa and aging (see Discussion).

Finally, to evaluate the gut microbiome’s potential to serve as a metagenomics-based “clock”, we developed a machine-learning model, aiming to predict a dog’s age based on its gut microbiome features. Specifically, we trained a random forest regression model on data from healthy dogs only, using 10-fold cross-validation. The model further incorporated information about the dogs’ sex, breed size, weight, and technical factors. Our model resulted in an average Pearson correlation coefficient of 0.350 between the real and predicted age of healthy dogs (Figure 3F). Importantly, to assure that the model’s ability to predict age was not driven solely by non-microbiome covariates, we compared the above model to a reference model that was trained using only the dog’s sex, breed size, weight, and technical factors, confirming that this reference model achieves a substantially lower correlation (Figure 3G; Pearson r=0.158).

### Clinical measures associated with the gut microbiome

Given the findings above and the potential role of the microbiome in promoting host health and in modulating various physiological and molecular host processes, we further examined the clinical laboratory data available for these dogs. To this end, we used complete blood counts (CBC) and chemistry profiles (CP) from peripheral blood, as well as urinalysis data, available for each member of the DAP Precision Cohort (n = 902 subjects). We first applied a mixed effects modeling approach, controlling for dogs’ age, general health status (owner-reported), sex, weight, breed size, diet, and technical covariates, to quantify how the microbiome and anthropometric measurements explain observed variation in the clinical urine and blood measurements (see Methods). We found that clinical measurements varied in their overall explained variance, ranging from 25.3% to 1.5% (Figure 4A). Notably, variation in platelet count (PLT), absolute lymphocyte count (abs Lymphocytes), and alkaline phosphatase (ALKP) exhibited the highest overall explainability among all measurements. We further examined the individual contribution of each microbiome or anthropometric factor to the overall explained variance. While the microbiome accounted for a modest portion of the variance, ranging from 0.4% to 3.47% (mean 1.61% ± 0.68%), it was in fact the most substantial factor in explaining variance across all lab tests, alongside age, and exceeded the influence of breed size, sex, health conditions, and weight (Figure 4B; FDR<0.05, two-sample Wilcoxon test).

**Figure 4:**
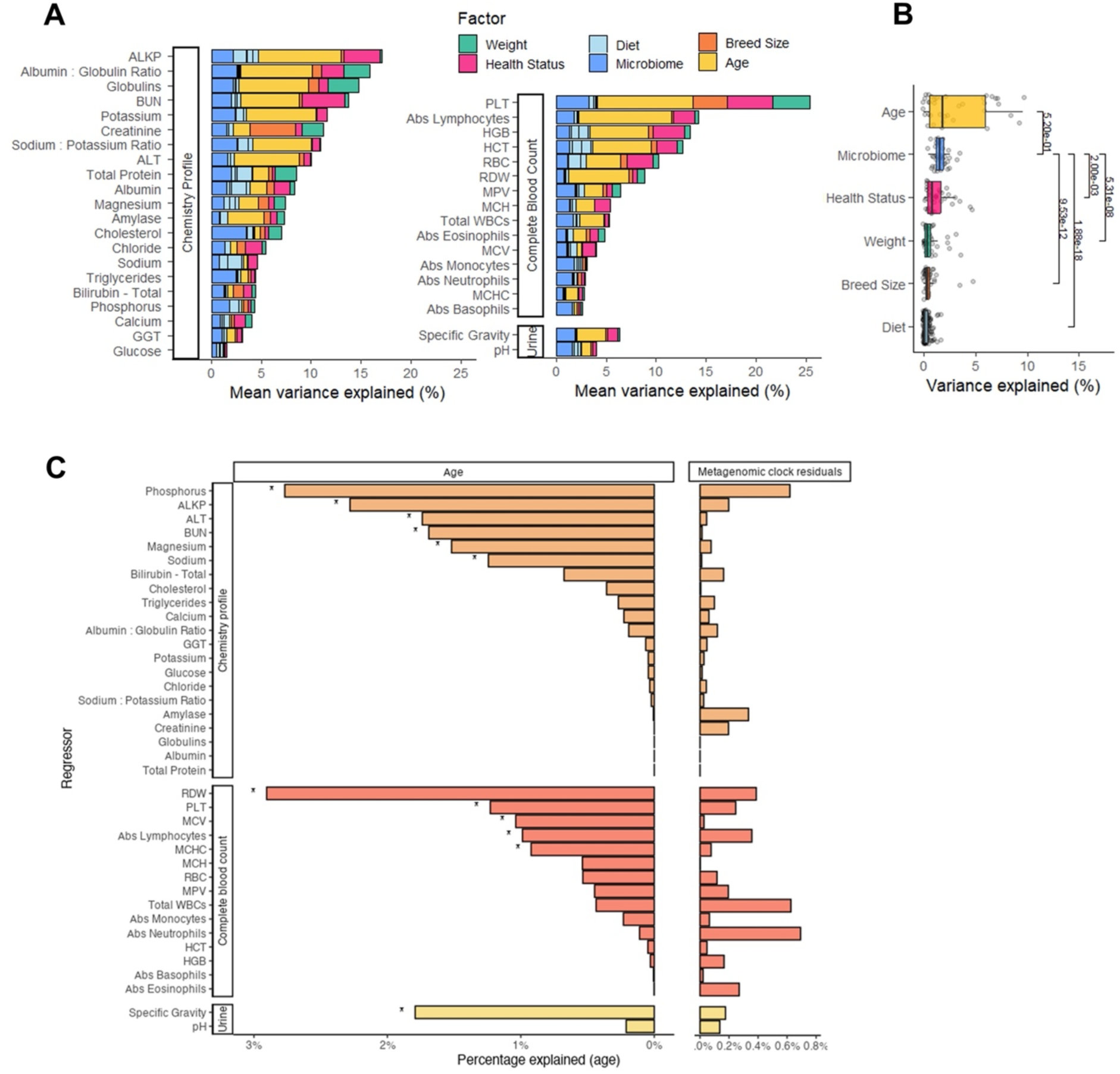
Association of the microbiome with clinical parameters. **(A)** Variation in blood lab test results explained by anthropometric measurements and the microbiome using fixed effects linear models. **(B)** The contribution of each factor to the variance explained across all clinical measurements. Significance of pairwise differences was computed with Wilcoxon tests and FDR corrected p-values are presented. **(C)** Variation in age explained by blood lab test results (left panel) and variation in metagenomic age residuals (right panel), assessed using permutation-based mixed effects models. Statistically significant results (FDR<0.05) are indicated with an asterisk. ALKP - Alkaline phosphatase, ALT - Alanine transaminase, BUN - Blood urea nitrogen, GGT - Gamma-glutamyl transferase, PLT - Platelet count, RDW - Red blood cell distribution width, Total WBC - total white blood count, HGB - Hemoglobin, HCT - Hematocrit, RBC - Red blood cell count, MPV - Mean platelet volume, MCH - Mean corpuscular hemoglobin, MCV - Mean corpuscular volume, MCHC - Mean corpuscular hemoglobin concentration.

Considering the strong association between the microbiome and clinical laboratory test results, we then characterized the pairwise correlation between each microbiome species or KEGG pathways and urine, CBC, and CP measurements (see Methods). In total, we identified 193 significant correlations (FDR<0.05), with 22 out of the 38 lab measurements exhibiting at least one significant correlation with either a microbiome species or a KEGG pathway (Supplementary Table S12). Notably, while many of these associations likely represent yet-uncharacterized mechanisms and are challenging to interpret, some can be explained by known physiological processes. For example, we found that blood cholesterol levels are positively correlated with microbial pathways involved in fatty acid and biotin synthesis (KEGG pathways ko01212 and ko00780). Indeed, an increase in biotin and fatty acid levels could elevate blood cholesterol, as biotin enhances fatty acid synthesis^37^, and may in turn increase production of acetyl-CoA, a key precursor in the cholesterol biosynthesis pathway. Similarly, we found an association between carbon metabolism and blood potassium levels, which is known to be mediated via insulin secretion^38^.

Finally, given this observed link between the microbiome and clinical parameters on the one hand, and the association between the microbiome and age reported above, we examined whether clinical parameters were associated with the residual difference between chronological age and predicted “metagenomic” age from our metagenomics clock predictions. While, as expected, we found that multiple clinical parameters were associated with chronological age (Figure 4C, left panel), no parameter exhibited a statistically significant contribution to the variance in metagenomics-based clock residuals (Figure 4C, right panel). This suggest that clinical parameters and the metagenomic clock capture different aspects of aging.

## Discussion

In this study, we present a comprehensive, population-wide analysis of the gut microbiome in companion dogs. Importantly, our study uniquely integrates analyses of associations between the canine microbiome and various lifestyle, environmental, and clinical factors, setting the DAP apart from other large-scale studies on the canine microbiome. Indeed, by sequencing stool metagenomes and coupling the obtained microbiome profiles with clinical parameters and owner-reported questionnaires of almost 1,000 dogs from across the United States, we identify key factors associated with the gut microbiome, including age, clinical measures, diet, and health. We specifically reveal significant age-associated shifts in the composition of the microbiome and introduce a metagenomics-based clock. Beyond the global association of the gut microbiome with the host diet, we also identify specific effects of the type of diet (commercial vs. home-prepared) and of coprophagy (feces eating) on the microbiome. We further highlight significant associations of the microbiome with multiple clinical blood and urine markers. Collectively, these insights deepen our understanding of the canine gut microbiome, the factors shaping it, and its relationship with dog health and age.

Importantly, our study offers a complementary viewpoint to microbiome studies performed in human population-wide cohorts. While host biology clearly differs significantly, humans and companion dogs share exposure to environmental factors, and it is thus interesting to assess differences and similarities in the factors driving microbiome variation. Indeed, some of our findings align with known determinants of the human gut microbiome. For example, aging in humans has been previously characterized by higher diversification with age^4^, and a similar trend was observed in our cross-sectional analysis of dogs presented here. Similarly, our findings concerning age-associated shifts in the abundances of *Prevotella* and *Holdemanella* agree with previous observations concerning these taxa and their putative role in aging. Specifically, *Holdemanella biformis* (previously known as *Eubacterium biforme*) is known for producing the long-chain fatty acid 3-hydroxyoctadecaenoic acid, which has an anti-inflammatory effect^39–42^. In line with our findings, literature reports a negative correlation between *Holdemanella biformis* and age, observed both in humans^43^ and in dogs^44^. Similarly, certain species within the *Prevotella* genus, such as *Prevotella copri*, have also been documented as decreasing in abundance with age in humans^5^ and in dogs^44^. However, *Prevotella* is also known for its substantial variability across dogs^45^, with some studies presenting contradictory evidence regarding its association with age^24,46^. These contradicting results highlight the need for careful consideration of cohort-specific effects, and the importance of large and population-wide cohorts. Moreover, both humans and canines show close association between clinical laboratory parameters and microbiome composition^47^. Finally, diet is identified as a key factor influencing microbial community structure, and processed commercial foods have been found to have a unique signature in both humans and dogs^48^. Hence, this work provides a gateway to study variation within the microbiome across host species.

We also recognize several potential sources of bias in this cohort that may influence our understanding of microbiome associations. First, age-related associations might be influenced by “survivor bias”, a common concern in prospective cohorts (such as the UK Biobank)^49^, as older dogs in the study may represent a healthier subset of the population, potentially due to inherent resilience or better access to healthcare. Additionally, as owners of dogs with specific health conditions or behavioral issues may be less inclined to enroll their pets in such a study, self-selection bias may be present^50^. With that in mind, the patterns and associations reported above may reflect primarily those of healthy aging. Furthermore, since the study was conducted in the USA, our findings may not accurately represent canine living conditions or microbiome diversity of dogs from other parts of the world.

This cross-sectional study establishes the baseline map of the canine microbiome, analyzing a single sample from each dog. Longitudinal data collection is ongoing and future data releases will enable a more dynamic characterization of molecular phenotypes, creating a unique opportunity to study interactions between aging and the microbiome in unprecedented detail. Moreover, DAP Precision Cohort dogs are characterized via complementary omics (including metabolomics, genomics, and epigenomics) providing an exciting vehicle to study the multi-omic signatures of aging. The Dog Aging Project has thus created a unique, citizen-science, and open-resource data repository, available for use by aging researchers, microbiologists, ecologists, epidemiologists, and veterinarians to elucidate diverse clinical and basic research questions.

## Methods

### DAP Precision 1 cohort description

The Dog Aging Project (DAP) is a multi-disciplinary prospective population-based cohort that aims to study aging in companion dogs^27^. The DAP *Precision* Cohort^29^ is a sub cohort within DAP that includes nearly 1,000 dogs for which multi-omic profiles were obtained, including stool metagenomics, blood metabolomics, whole genome sequencing, and epigenetics. Clinical lab tests, including complete blood counts (CBC), blood chemistry profile (CP) and urine analysis, were additionally collected. Lastly, dog owners were required to complete multiple surveys about their dogs’ health status, diet, living environment, and lifestyle, both at the time of enrollment and yearly thereafter. In this study, we analyzed only the baseline sample from each DAP Precision Cohort dog.

### Stool specimen collection, extraction, and sequencing

Stool samples were collected by primary care veterinarians using PB-200 (DNAgenotek, Ontario, Canada) tube with agitator, and shipped on dry ice for processing and sequencing at the Snyder-Mackler Lab. Upon arrival, the samples were inventoried at −80°C until extraction. For each fecal sample, a lysate was prepared using the following procedure: The fecal collection tube was thawed and vortexed to homogenize the fecal sample in the storage solution. A small aliquot (∼200 µL) of the fecal slurry was removed using a fresh 1 mL serological pipette and transferred to a 2 mL tube pre-filled with garnet shards and a 6 mm zirconium bead (Benchmark Scientific, D1033-30G). To this tube, 365 µL of PowerBead solution (Qiagen, #12955-4-bs) and 20 µL of 20% sodium dodecyl sulfate (SDS) were added. The tubes were subjected to bead-beating using a TissueLyzer instrument for 45 seconds at 30 cycles per second. The resulting fecal homogenate was treated twice with Protein Precipitation Solution (MP Biomedical, product no. 116560203) to remove proteins. The fecal lysate was then inventoried and stored in 2 mL tubes at −80°C.

Metagenomic libraries were generated from fecal lysates or extracted DNA using a published protocol. Briefly, libraries are constructed using the Illumina DNA Prep kit (20060060) adapted for half-reactions. Libraries were quantified using a plate reader and QuantSeq (Invitrogen, Waltham, MA) fluorescent dye. Libraries were pooled in equimolar amounts into pools of 120 samples. Pools were run through a Pippin Prep instrument (Sage Science, Beverly, MA) to remove fragments <350bp. Metagenomic library pools were sequenced on the Illumina NovaSeq platform, with a goal of > 5 million reads/library.

### Metagenomic data processing

First, paired-end reads were filtered to remove low-quality reads, short reads (below 60bp) and duplicated reads using fastp^51^ (version 0.23.2; Executed with the ‘dedup’ and ‘length_required’ flags). Next, host-associated reads, as determined by alignment to a reference canine genome (CANFAM4^52^) with BOWTIE2^53^ (version 2.3.5), were removed. The remaining reads were rarefied to 5M reads per sample, and samples with less than 5M reads were discarded. After the above pre-processing, 922 samples remained. The average *raw* sequencing depth of these 922 samples was 48M reads (sd: 45.2M). Next, taxonomic profiles according to the GTDB taxonomy^32^ (release 207) were produced using Kraken2^54^ (version 2.1.1) and relative abundances at the phylum, genus and species level were estimated using Bracken^55^ (version 2.8). Rare taxa that appeared in <20% of the samples, or with an average relative abundance <0.0001, were removed. In addition, functional microbiome profiles based on KEGG pathways were estimated using MetaLaffa^56^ (version 1.0.1).

### Lab test and genetic data

Blood samples were collected in EDTA tubes, from either one draw of 20 mL for dogs >8 kg, or 12 mL drawn six weeks apart for dogs ≤8 kg. Tubes were shipped to the Texas Veterinary Medical Diagnostic Laboratory. Complete Blood Counts (CBC) were performed on Advia 120 Hematology System (Siemens Medical Solutions, Malvern, PA). Blood chemistry profiles were obtained from serum extracted from the additive-free tube included in DAP biospecimen kits, using the DxC700AU Chemistry Analyzer (Beckman Coulter, Brea, CA).

Urinalysis was performed with approximately 3 mL of urine. Urine specific gravity was calculated by refractometer; chemical analysis was performed with Multistix 10 SG Urine Test Strips (Siemens Medical Solutions, Malvern, PA); and microscopy was performed manually.

DNA kits with barcoded cheek swabs for collecting cheek samples were shipped to dog-owners (GBF, Inc., High Point, NC). Swabs were mailed to Neogen/GeneSeek where low-pass DNA sequencing was performed.

### Comparison of DAP to Coehlo et al

We compared the gut microbiomes of DAP samples to those from another shotgun-sequenced dog cohort described in Coehlo *et al.*^18^. Briefly, the authors of that study performed a diet intervention study in 64 dogs (32 Labrador retrievers and 32 beagles), collecting stool samples once at baseline and again following intervention. We obtained raw shotgun sequencing data of baseline samples only from the European Nucleotide Archive (study accession: PRJEB20308), and dog metadata from the publication’s supplementary information. Raw sequencing data was processed into taxonomic profiles using the same processing pipeline described above. Average taxon abundances were compared between the two cohorts using Spearman correlations, at varying taxonomic levels.

### Comparison of DAP to Branck et al

We compared the taxonomic profiles of DAP dogs, obtained by the pipeline above, to taxonomic profiles obtained in another dog study that was recently published and that was based on a different profiling approach^20^. Using supplementary data from that study, we computed the average relative abundances of each species in the 1,187 household dogs included in their cohort. Of the top 100 abundant species, only 44 were mappable with high confidence to the GTDB taxonomy used here. For this subset of species, we calculated the Spearman correlation between the average relative abundances in the two cohorts.

### Statistical analysis

#### Covariation between the microbiome and other data types

To estimate the global association between the gut microbiome’s taxonomic and functional profiles and other available data about the dogs’ lifestyle, health, diet, etc. (“data types”), we first computed between-dog distance matrices per data type (see below). Using these distance matrices, we then quantified the covariation between each pair of data types using either Mantel tests or Procrustes’ permutation test. P-values were corrected for multiple hypothesis testing using the Benjamini–Hochberg procedure (FDR). We used the ‘mantel’ and ‘protest’ functions, respectively, from the ‘vegan’ package (version 2.6-2)^57^. The number of permutations for both tests was set to 999.

Distance matrices were calculated as follows: Microbiome-related distance matrices were based on Bray-Curtis, applied to species relative abundance data. Genotype distances were based on the genetic relationship matrix calculated using PLINK2^58^. For the ‘Sex’ distance matrix, distances between dogs were defined as either 0 if two dogs shared the same sex category (4 categories accounting for sterilization status), and 1 otherwise. For all other data types, Euclidean distances were used. In Supplementary Table S2, we list for each data type the exact list of features that were included, and the final number of Precision dogs for which all data was available.

As Mantel and Procrustes both examine the association between each pair of distance matrices independently, ignoring possible dependencies between data types, we also used ‘Multiple regression on distance matrices” (MRM) as implemented by the “MRM” function from the “ecodist” R package (version 2.0.9)^59^. Here, we tested the association between Bray-Curtis microbiome distance matrices and each data type, while controlling for all other data types. All Mantel, Procrustes, and MRM results are provided in Supplementary Tables S3 and S4.

### Predicting abundances of specific genera using various data types

We trained random forest regression models for each of the top 20 most abundant genera (Supplementary Table S1), using each of the different data types, and evaluating the models using repeated 5-fold cross-validation (10 repeats). We summarized each model’s performance by averaging the Spearman correlation coefficients between predicted and actual abundance values, over all folds and repeats. Random forest models were trained using the “ranger” R package (version 0.14.1) and using default parameters.

### Associations between specific microbial taxa or functions, and host phenotypes

To identify associations between specific taxonomic or functional features of the microbiome and various host phenotypes, we used mixed effects linear models, as implemented in the “MaAsLin2” R package^60^ (v.1.12.0). Specifically, the relative abundance of the microbiome feature (either a taxon or a pathway) was modeled as a function of the specific host phenotype, as well as the dog’s health status, breed size (standard breed size groupings), weight, sex, metagenomic sequencing depth and duplication rate as fixed effects. Additionally, sequencing batch was included as a random effect. P-values were FDR corrected.

### Associations between overall microbiome composition and host age

We employed PERMANOVA^61^ (Permutational Multivariate Analysis of Variance) to assess global differences in microbial community composition across four age groups, also referred to as “life stages”. These groups were defined based on the American Animal Hospital Association guidelines (https://www.aaha.org/resources/life-stage-canine-2019/) and were adjusted for different breed sizes. Using the adonis2 function from the “vegan” R package (v.2.6.4), we incorporated fixed effects as described above. Sequencing batch was included as another fixed effect, as adonis2 does not support random effects. We ran the function with default parameters and 999 permutations.

### Microbiome uniqueness score

We calculated the uniqueness score for each dog by determining the distance between each sample to its nearest neighbor, using the Bray-Curtis dissimilarity matrix, and following the methodology described in Wilmanski *et al.*^4^. This approach allowed us to quantify how distinct each dog’s gut microbiome was in comparison to other dogs in the cohort.

### Age prediction model

We employed a random forest model using the R ranger package (v.0.16.0) to predict dogs’ ages based on their gut microbiome taxonomic profiles at the species level, incorporating the dog’s sex, breed size, weight, and technical characteristics such as batch, metagenomic sequencing depth, and duplication rate. The model was trained on data from 843 dogs reported by their owners as having ‘excellent’ or ‘very good’ health. We performed 10-fold cross-validation, predicting ages in each fold to evaluate the model’s performance and minimize overfitting. Pearson correlation coefficients between predicted and actual ages were calculated for each fold.

Additionally, we trained another model using only the dog’s sex, breed size, weight, and technical characteristics (excluding microbiome data) to confirm that the predictive signal achieved by the previous model was microbiome-driven.

### Lab test analysis

To estimate the extent to which different factors explain the variance in each clinical blood and urine measurement, we employed a linear modeling approach. In each model we included the dog’s age, weight, breed size, health status, microbiome composition, diet diversity, and sequencing quality metrics (as previously noted). Given the high dimensionality of microbiome data, we used the first 10 principal coordinates (PCoA) from the Bray-Curtis distance matrix, and for diet diversity, we used the first four principal components (PCs). To account for possible effects of feature orderings in each model, we applied a permutation-based approach as recommended by Grömping^62^. To compare differences in the variance explained by different features, we conducted pairwise Wilcoxon tests with a final FDR correction.

### Clinical blood measurements and age variation

To estimate how well each blood measurement predictor explains the variance in age, we calculated Eta-squared (η²) using ANOVA Type II with the “effectsize” R package (v0.8.9)^63^. This method quantifies the proportion of age variance attributable to each blood measurement predictor while accounting for all other predictors in the model, estimating their relative contributions.

To explore metagenomic changes associated with age, we performed linear regression with real age as the predictor and metagenomic predicted age (from the metagenomics-based clock) as the response. We analyzed the residuals—differences between observed and predicted ages— to identify metagenomic changes not explained by chronological age. We then assessed the variance in these residuals attributable to each blood measurement predictor, using the same approach as for the age variance analysis.

## Supplementary Figures

**Supplementary Figure S1:**
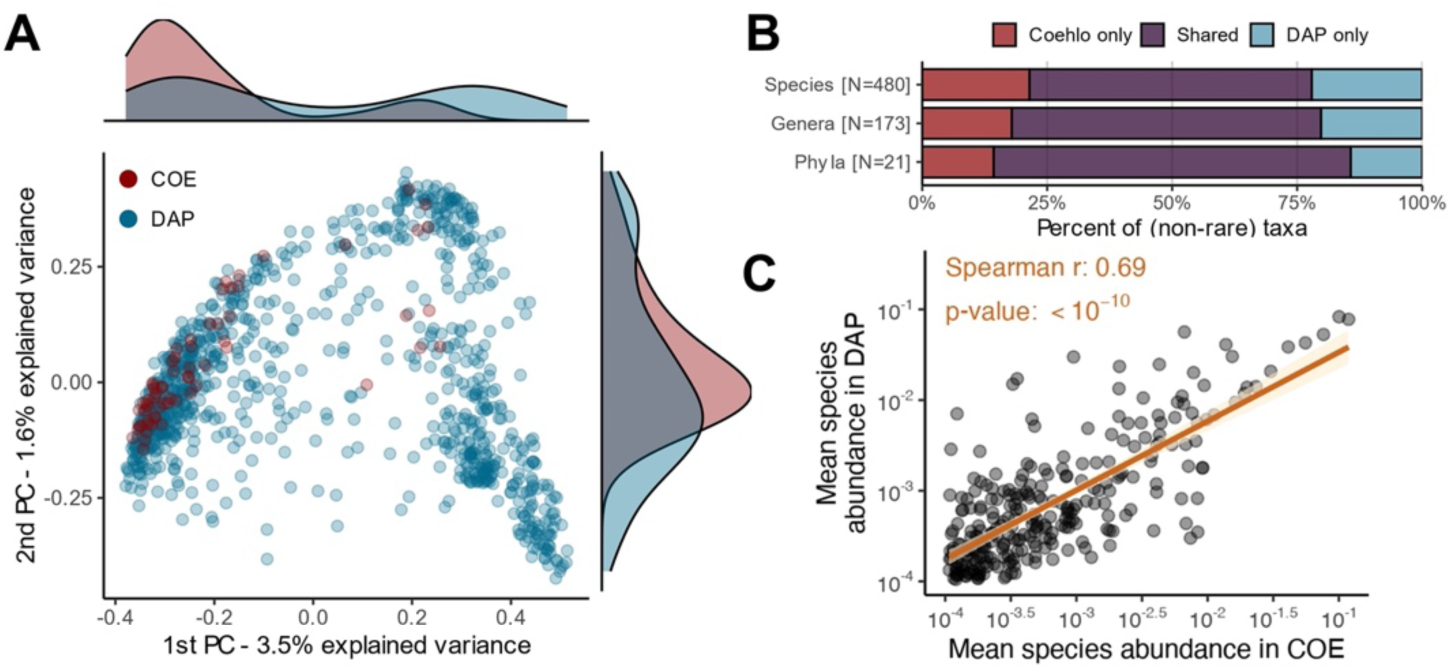
Comparison between gut microbiome taxonomic profiles of the DAP cohort and Coehlo *et al.* cohort. **(A)** Principal coordinates analysis (PCoA) based on species-level Bray-Curtis dissimilarity matrices, including both the DAP cohort and the cohort from Coehlo *et al.* (COE). **(B)** Percent of species, genera and phyla, that were identified in both cohorts or in only one of the cohorts. In each cohort, only taxa that appeared in >20% of the samples, with an average relative abundance >0.0001 were considered (i.e., “non-rare”). **(C)** A comparison of average relative abundance of each species, between the two cohorts. See Figure 2B.

**Supplementary Figure S2:**
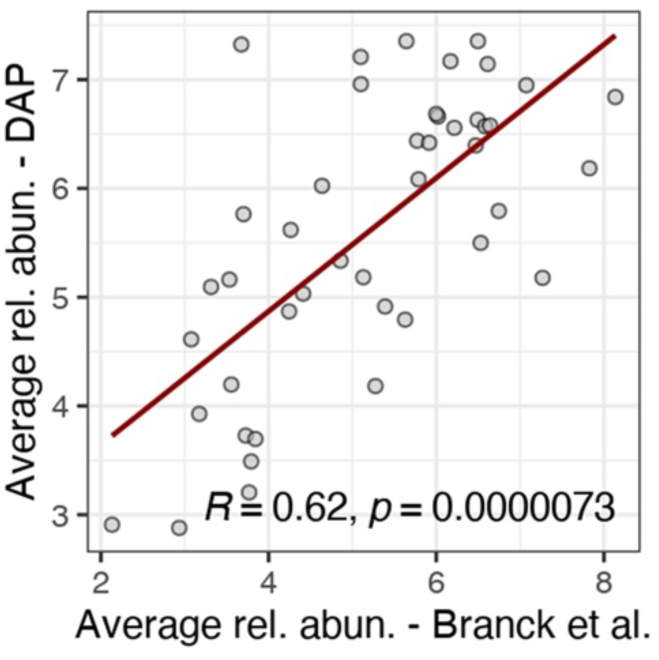
Comparison between gut microbiome taxonomic profiles of the DAP cohort and Branck *et al.* cohort. A comparison between the average relative abundances of gut species in the DAP cohort, computed using a reference-based approach, and gut species reported in Branck et al., computed using a combined assembly- and reference-based approach.

**Supplementary Figure S3:**
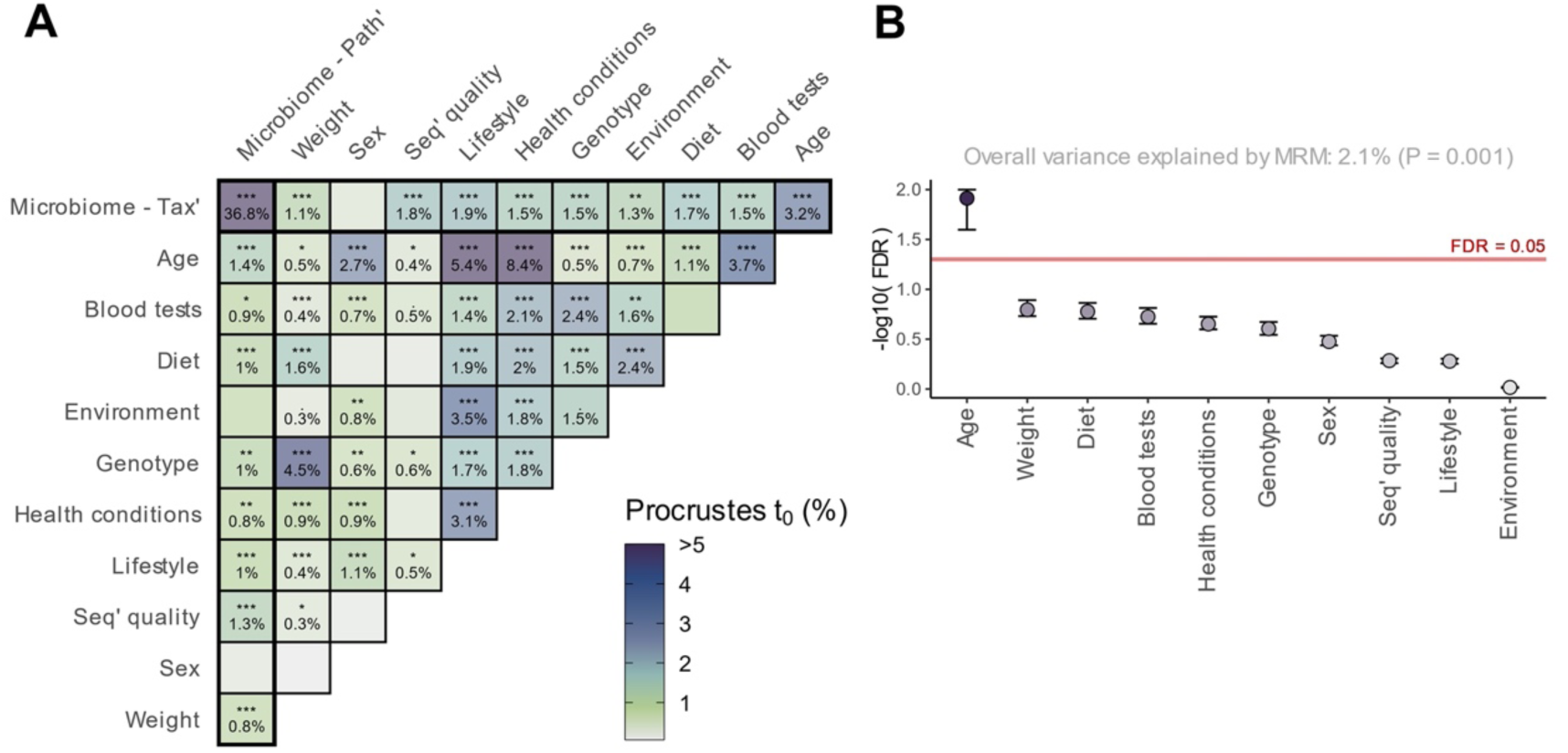
Broad associations between taxonomic and functional profiles of the dog gut microbiome and multiple demographic, lifestyle, and other factors. **(A)** Pairwise Procrustes tests to estimate the associations between each pair of measurement types. Tiles are colored by the Procrustes statistic (Spearman’s rho), roughly equivalent to shared variance. Stars indicate FDR-corrected p-value significance (‘.’ <= 0.1 ‘*’ <= 0.05, ‘**’ <= 0.01, ‘***’ <= 0.001). See Supplementary Table S3. Tax’: Taxonomy, Path’: Pathways, Seq’: Sequencing. **(B)** Coefficient p-values from a multiple regression on distance matrices (MRM), predicting microbiome species-level Bray-Curtis distances from multiple distance matrices based on various demographic, lifestyle, and other dog factors.

